# Hippocampus-striatum wiring diagram revealed by directed stepwise polysynaptic tracing

**DOI:** 10.1101/2021.10.12.464132

**Authors:** Wenqin Rita Du, Elizabeth Li, Jun Guo, Yuh-tarng Chen, So Jung Oh, Aspen Samuel, Ying Li, Hassana K Oyibo, Wei Xu

## Abstract

The hippocampus and the striatum represent two major systems in the brain for learning, memory and navigation. Although they were traditionally considered as two parallel systems responsible for distinct types of learning or navigation, increasing evidence indicates a close synergetic or competitive interaction between them. Both the hippocampus and the striatum consist of multiple anatomical and functional domains. Besides the limited direct projection from the hippocampus to the ventral striatum, most of the functional interaction between them may be mediated by polysynaptic projections. Polysynaptic connectivity has been difficult to examine due to a lack of methods to continuously track the pathways in a controlled manner. Here we developed a novel approach for directed stepwise polysynaptic tracing by reconstituting a replication-deficient retrograde transneuronal virus – pseudorabies virus lacking gene IE180 (PRV^ΔIE^). We minimized PRV neurotoxicity by temporally restricting viral replication; and enabled both anatomical tracing and functional analysis of the circuits. With these tools, we delineated a hippocampus-striatum wiring diagram, which consists of pathways from specific functional domains in the hippocampus to corresponding domains in the striatum via distinct intermediate regions. This polysynaptic wring diagram provides a structural foundation for further elucidation of the interaction between the hippocampus and the stratum in multiple brain functions.

**Significance Statement:** We created a new method for controlled stepwise tracing of polysynaptic neuronal circuits which also enables functional analysis of the circuits. With this method, we revealed the polysynaptic wiring diagram between the hippocampus and the striatum, two major brain structures centrally involved in learning, memory and spatial navigation. This wiring diagram demonstrates how specific anatomical domains in the hippocampus are connected to the specific domains in the striatum via distinct intermediate brain regions; thus it will help to elucidate the functional interaction between these two major memory and navigation systems. Our new method can be broadly applied to many other brain circuits for anatomical and functional analysis.

## Introduction

The hippocampus and the striatum are two major brain regions involved in learning and navigation. They are frequently considered as two independent systems for distinct types of learning or navigation. The hippocampus is critical for explicit/declarative memories encoding facts and events, which can be formed through a single trial (Buzsaki and Moser, 2013). The striatum is critical for stimulus-response association and implicit memories including procedure memories, which normally requires repetitive practice (Graybiel and Grafton, 2015). Furthermore, the striatum is preferentially involved in egocentric navigation in which animals rely on body direction and local cues for navigation; while the hippocampus preferentially contributes to allocentric navigation in which animals navigate with a world-centered spatial “map” of the environment (Chersi and Burgess, 2015).

Despite the functional division, accumulating evidence indicates a close interaction between the hippocampus and the striatum. Firstly, the hippocampus is involved in acquiring and consolidating procedural memory and motor skills. Human imaging studies demonstrate highly correlated activations of the striatum and hippocampus in learning novel motor sequences (Fernandez-Seara et al., 2009; Gheysen et al., 2010), and the contribution of the hippocampus to consolidating motor memories (Albouy et al., 2013). Secondly, the striatum contributes to declarative memory, especially the retrieval of the declarative memory (Scimeca and Badre, 2012). Thirdly, the hippocampus and the striatum collaborate and compete in spatial navigation (Goodroe et al., 2018). The close functional relationship between the hippocampus and striatum is also supported by the coupling of the hippocampal and the striatal neuronal activities (Lansink et al., 2016; Sjulson et al., 2018; Tort et al., 2008).

Both the hippocampus and the striatum contain multiple anatomical sub-regions/domains with distinct functions (Bienkowski et al., 2018; Fanselow and Dong, 2010; Voorn et al., 2004). Mapping the anatomical wiring diagram between the anatomical domains is essential to understanding hippocampus-striatum interaction. The hippocampus and striatum are not directly connected except that nucleus accumbens (NAc) in the ventral striatum receives inputs from the hippocampus, mainly the ventral hippocampus (Brog et al., 1993; Groenewegen et al., 1987; Kelley and Domesick, 1982; McGeorge and Faull, 1989; Trouche et al., 2019). Their functional interaction can therefore be mediated through polysynaptic pathways.

However, the polysynaptic pathways are technically challenging to delineate. A few neurotropic viruses, such as pseudorabies virus (PRV), vesicular stomatitis virus (VSV), rabies virus (RV) and herpes simplex virus (HSV), can replicate inside neurons and spread across synapses. This property makes them self-amplifying tracers for delineating polysynaptic pathways (Astic et al., 1993; Blessing et al., 1994; Card et al., 1991; Lundh, 1990). However, these replication-competent viruses have significant limitations. Firstly, the viruses are highly toxic, making them unsuitable for functional analysis (Enquist, 2002). Secondly, the viruses spread to all connected brain regions and cannot be selectively directed to a particular pathway. Frequently, a large number of brain regions become infected after 2-3 orders of transneuronal spreading, making it difficult to identify the exact routes the viruses traveled on. Thirdly, the viruses may spend different amounts of time spreading across one order of connection depending on the length of axons or other neuronal properties, making it hard to determine the sequence of synaptic connections.

Here we first created an inducible, directed and stepwise transneuronal tracing system by inducibly reconstituting a replication-deficient PRV, which lacks immediate gene IE180 (Oyibo et al., 2014). In this new system, transneuronal spreading of PRV can be activated 1) at a desired time, 2) at a selected branch of circuit, and 3) to pass a pre-determined number of orders of synaptic connections. The replication of PRV is restricted to a short time window to avoid potential neurotoxicity, enabling functional analysis. We then used these tools to delineate the wiring of the hippocampus-striatum pathways. We found that the different functional domains of the hippocampus are selectively wired to the different regions in the striatum via distinct intermediate regions.

## Materials and Methods

### Mice

6-7 weeks old male C57BL/6J mice (UT Southwestern breeding core, JAX), or tdTomato reporter mice, the Ai9 mice (JAX stock No. 007909), were group housed on a 12 hr light /12 hr dark cycle with ad libitum access to food and water. Mice were randomly assigned to experimental groups. Animal work was approved and conducted under the oversight of the UT Southwestern Institutional Animal Care and Use Committee and complied with Guide for the Care and Use of Laboratory Animals by National Research Council.

### Cell culture

HEK293 cells, 3T3 cells and Pk15-IE180 cells were used to generate AAV, test PRV replication and amplification of PRV^ΔIE^, respectively. HEK 293 cells and 3T3 cells were grown in Dulbecco’s Modified Eagle’s Medium (DMEM) containing 10% FBS. Pk15-IE180 cells were grown in DMEM containing 10% FBS and 700 μg/mL of G-418.

### Viral vector construction

AAV vectors were constructed with AAV2 inverted terminal repeats (ITRs). In AAV-tTA or AAV-rtTA, the following components were arranged sequentially downstream of left-ITR: synapsin promoter, tTA or rtTA, Posttranscriptional Regulatory Element (WPRE), hGH poly A sequence, and the right ITR. In AAV-IE180WT and AAV-IEo, the following components were arranged sequentially downstream of left-ITR: the tetracycline-responsive element and a minimal CMV promoter, the coding sequence of wild type IE180 or IEo, SV40 poly A sequence, and the right ITR. In SynaptoTAG2 AAV the following components were arranged sequentially downstream of left-ITR: synapsin promoter, tdTomato, P2A, EGFP fused with Syb2, WPRE, hGH poly A sequence, and the right ITR. In AAV-NS, the following components were arranged sequentially downstream of left-ITR: synapsin promoter, NS1, P2A, dTomato with SV40 nuclear localization signal or sequence (NLS), WPRE, hGH poly A sequence, and the right ITR. In AAV-dfi-NS1, the following components were arranged sequentially downstream of left-ITR: synapsin promoter, loxp site and lox2272 site, inverted open reading frame consisting the coding sequences of NS1, P2A and NLS-dTomato, Loxp site, Lox2272 site, WPRE, hGH poly A sequence, and the right ITR. Besides the AAV vectors constructed in this study, the AAV mediating Cre-dependent expression of hM4Di was requested from Addgene (44362, Krashes MJ et al., 2011).

### Virus production

All AAVs except rAAV2-retro-cre were packaged with AAV-DJ capsids. AAV2-retro-cre was packaged with rAAV2-retro capsid genes (rAAV2-retro helper plasmid). Virus was prepared as described (Zolotukhin et al., 1999). Briefly, AAV vectors were co-transfected with pHelper and pRC-DJ or rAAV2-retro helper into AAV-293 cells. Cells were collected 72 hr later, lysed, and loaded onto iodixanol gradient for centrifugation at 400,000 g for 2 hr. The fraction with 40% iodixanol of the gradient was collected, washed, and concentrated with 100,000 MWCO tube filter. The genomic titer of virus was measured with quantitative real-time PCR. The titers of AAVs used for stereotaxic injection were in the range of 0.5-2 × 10^13^ copies/ml.

IE180-null PRV was amplified by infecting Pk15-IE180 cells, which express IE180 in a Dox-dependent manner, as described (Oyibo et al., 2014). Briefly, Pk15-IE180 cells were infected with a stock of IE180-null PRV at the presence of 1 ug/ml of doxycycline (MP Biomedical, 219895501). 3-4 days later, the infected cells, after showing apparent cytopathic effect, were lysed by three freeze-thaw cycles and sonication. _Cell lysate was clarified by centrifuge for 10 min at 3_000 g. _The supernatant was laid on top of a sorbitol buffer and centrifuged for 2 hr at 4°C at 23.5k rpm with a SW28 rotor. The pellet was resuspended in DMEM. Besides_ IE180-null PRV, the replication-capable PRV--PRV154--was from CNNV core.

### Stereotaxic viral injection

Mice were anesthetized with tribromoethanol (125–250 mg/kg). Viral solution was injected with a glass pipette at a flow rate of 0.10 μl/min. After the completion of injection, the glass pipette was left in place for 5 min before being retrieved slowly. The coordinates for each of the injection sites on the anterior-posterior direction from the bregma (AP), medial-lateral direction from the midline, and the dorsal-ventral direction from the dura (DV) can be found in the following table. On the AP direction, “+” denotes anterior to the bregma and “-“ denotes posterior to bregma. We injected between 0.25 – 0.5 μl of viral solution at each injection site unless stated otherwise. The injections were unilateral except those injected for behavioral tests.

| Region | dorsal CA1 | ventral CA1 | dorsal CA3 | DG | dSUB | LS | PFC |
| --- | --- | --- | --- | --- | --- | --- | --- |
| **AP** | -1.95 | -3.00 | -2.25 | -2.00 | -3.15 | +0.55 | +1.25 |
| **ML** | 1.35 | 3.50 | 2.75 | 1.25 | 2.00 | 0.45 | 0.30 |
| **DV** | 1.20 | 5.00 | 2.50 | 2.20 | 1.75 | 2.70 | 1.25 |
| Region | dorsal striatum | NAc | rRSP | vcRSP | SNr | SUM | EC |
| **AP** | +0.55 | +1.00 | -2.25 | -3.40 | -3.15 | -2.50 | -4.80 |
| **ML** | 1.65 | 1.10 | 0.25 | 0.75 | 1.15 | 0.00 | 3.60 |
| **DV** | 3.00 | 5.20 | 0.75 | 1.25 | 5.50 | 5.75 | 2.60 |

### Histology and microscopy

Mice were transcardially perfused with 10 ml of PBS followed by 40 ml of 4% paraformaldehyde (PFA) in PBS. The brains were extracted and postfixed overnight in 4% PFA at 4°C, and cryoprotected in 30% sucrose. Brains were sectioned with a cryostat to a thickness of 40-μm. Free-floating sections were washed in PBS, mounted on glass slides and sealed with VECTASHIELD antifade mounting medium (Vector Laboratories, CA, Cat# H-1200). The whole-mount brain sections were scanned with Zeiss AxioscanZ1 digital slide scanner with a 10X objective. The high-resolution images were taken with ZEISS LSM 880 with Airyscan confocal microscope.

### Brain slice electrophysiology recording

3 weeks after PRV^ΔIE^--cre injection in CA1, transverse slices of the dorsal hippocampus (with a thickness of 300 μm) were prepared with a vibratome (Leica VT1200) in ice cold cutting solution containing (in mM): 2.5 KCl, 1.2 NaH_2_PO_4_, 26 NaHCO_3_, 10 D-glucose, 213 sucrose, 5 MgCl_2_, 0.5 CaCl_2_. The slices were incubated in cutting solution for 30 min at 32 °C and then in artificial cerebrospinal fluid (ACSF) containing (in mM): 124 NaCl, 5 KCl, 1.2 NaH_2_PO_4_, 26 NaHCO_3_, 10 D-glucose, 1 MgCl_2_, 2 CaCl_2_ for at least 1 h at 32 °C. The cutting solution and ACSF were adjusted to pH 7.3-7.4 and 290 - 300 mOsm and constantly aerated with 95% O_2_/5% CO_2_. Whole-cell patch clamp recording was performed in a recording chamber perfused (~1 ml/min) with oxygenated ACSF at 26-28 °C. The recording pipettes (2-4 MΩ) were filled with internal solution containing (in mM): 125 K-gluconate, 20 KCl, 4 Mg-ATP, 0.3 Na-GTP, 10 Na_2_-phosphocreatine, 0.5 EGTA, 10 HEPES, adjusted to pH 7.3-7.4 and 310 mOsm. Post-synaptic currents were recorded in voltage-clamp mode with holding potential at −70 mv. The resting potentials and current injection-induced action potentials were recorded in current-clamp mode.

## Experimental Design and Statistical Analysis

Data are presented as mean ± standard error of the mean (SEM). Sample number (n) indicates the number of cells or mice in each experiment and is specified in the figure legends. The data in Fig. 2 were analyzed with two-tailed Student’s t-test. p < 0.05 is considered statistically significant.

## Results

### 1) Reconstitution of PRV replication and transsynaptic transport with optimized IE180 (IEo)

IE180-null PRV, referred to as PRV^ΔIE^, can infect neurons but no longer replicate inside neurons or spread across synapses (Oyibo et al., 2014). To reconstitute the replication of PRV^ΔIE^ in the brain, we first made AAV constructs mediating the expression of wild type IE180 gene, but they did not produce reliable reconstitution (Fig. 1). IE180 has a GC-rich sequence (80.2% GC, whose function is not clear). So we modified IE180 coding sequence by lowering its GC content to 61.8% and named it IEo (o: codon-optimized).

**Figure 1.**
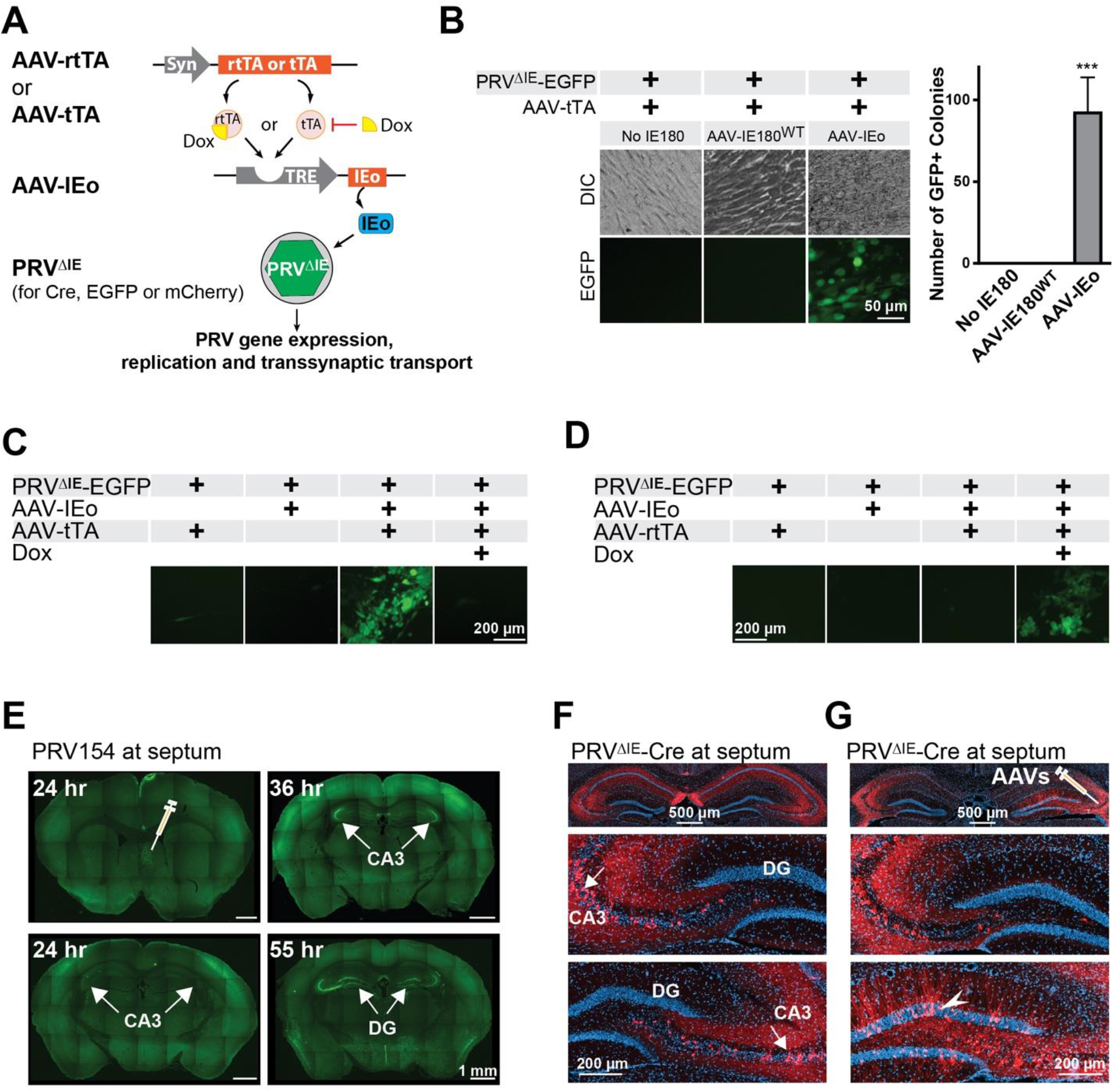
Reconstitution of PRV replication and transsynaptic transport with IEo. (**A**) AAV vectors (AAV-IEo and AAV-tTA or AAV–rtTA) were generated for inducible expression of codon-optimized IE180 gene (IEo) to activate the replication of replication-deficient PRV (PRV^ΔIE^). (**B**) Representative photos and quantification showing that the combination of AAV-tTA and AAV-IEo, but not that of AAV-tTA and AAV-IE180WT (AAV mediating expression of wild-type IE180), efficiently activated PRV^ΔIE^-EGFP gene expression and viral replication in cultured 3T3 cells (EGFP-expression was quantified by counting GFP+ colonies in each well of 6-well culture plates. The numbers from each batch of experiments were averaged. n=4 independent experiments.). (**C, D**) AAV-IEo and AAV-tTA (**C**) or AAV-IEo and AAV-rtTA (**D**) activated PRV^ΔIE^ gene expression and viral replication in cultured 3T3 cells in the absence or presence of doxycycline (Dox), respectively. (**E**) Replication-capable PRV expressing EGFP (PRV154) was injected into the septum unilaterally. EGFP-positive neurons were detected sequentially in the injection site, bilateral CA3 and bilateral dentate gyrus (DG) 24, 36 and 55 hours after viral injections, respectively. (**F**) PRV^ΔIE^ expressing cre (PRV^ΔIE^-Cre) was unilaterally injected into the septum of tdTomato reporter mice (Ai9 mice). tdTomato-positive neurons were detected at the bilateral CA3 (top panel, and enlarged photos in the lower 2 panels). No tdTomato-positive neurons were detected in DG contralateral (enlarged photo in the middle panel) or ipsilateral (lower panel) to the septal injection. (**G**) AAV-IEo and AAV-tTA were co-injected into CA3 unilaterally followed by an injection of PRV^ΔIE^-Cre in the septum ipsilateral to AAV injection. tdTomato-positive neurons were detected in the bilateral CA3 and in DG ipsilateral to AAV injections (lower panel) but not in contralateral DG (middle panel).

**Figure 2.**
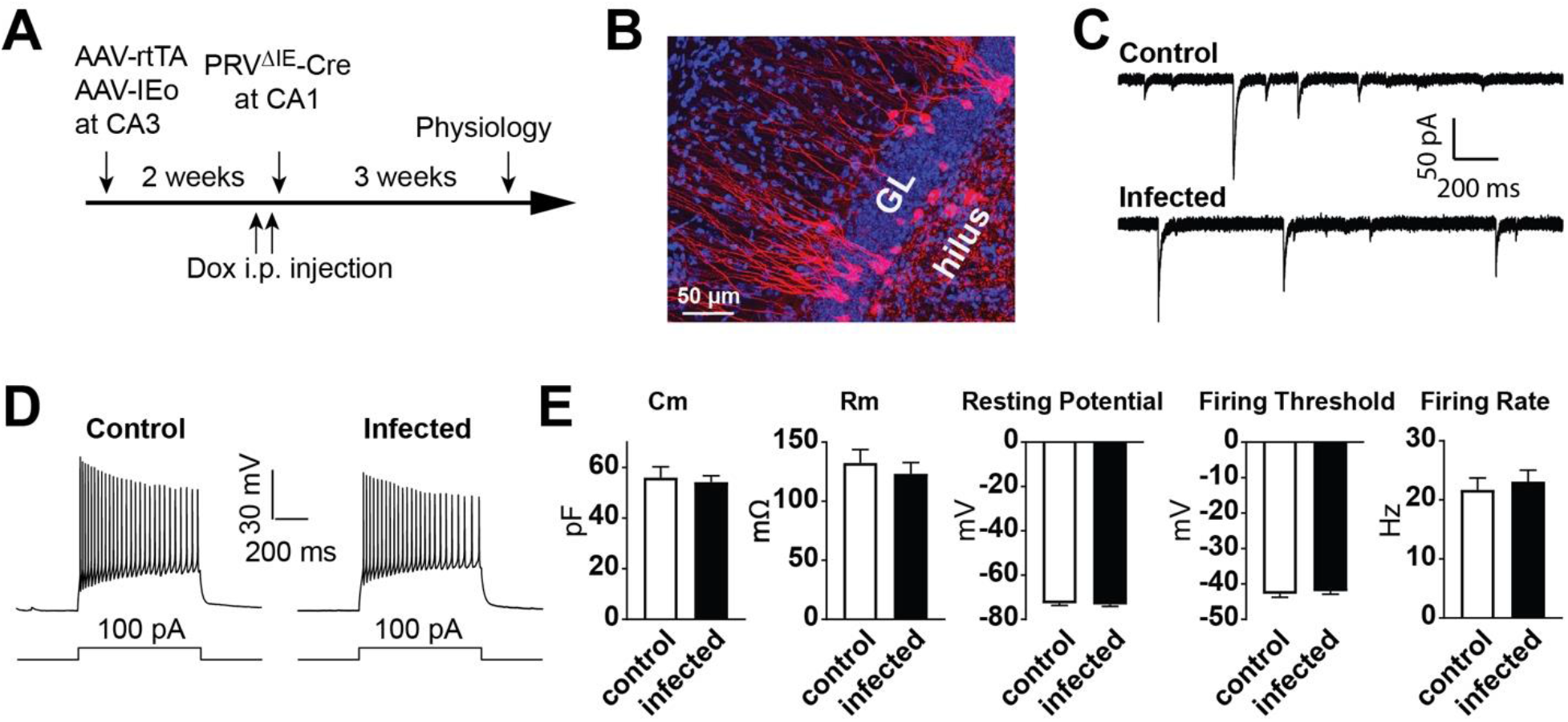
Temporally restricted replication minimized PRV neuronal toxicity. (**A**) Time course of electrophysiological analysis of PRV^ΔIE^-infected neurons in Ai9 mice. (**B**) Morphology of tdTomato-positive neurons in DG. (**C, D**) Whole-cell patch clamp recordings of tdTomato-positive neurons and their neighboring non-infected control neurons. Both control and PRV^ΔIE^-Cre-infected neurons demonstrated spontaneous synaptic currents (**C**) and action potentials triggered by current injections (**D**). (**E**) Quantification of electrophysiological parameters recorded from control and PRV^ΔIE^-Cre-infected neurons respectively. Cm: membrane capacitance; Rm: membrane resistance. Data are represented as mean ± SEM. n=14-15 neurons from 3 mice. No statistical significant difference was found between the two groups (Student’s t-test).

We constructed two sets of AAV vectors for inducible expression of IEo: one set (including AAV-tTA and AAV-rtTA) expresses the tetracycline-controlled transactivator (tTA) or the reverse tetracycline-controlled transactivator (rtTA) under the control of synapsin promoter; and the other (AAV-IEo) expresses IEo under the control of a promoter containing tetracycline-responsive element (TRE) (Fig. 1A). tTA activates IEo expression in the absence of doxycycline (Dox), while rtTA induces IEo in the presence of Dox. IEo, once expressed, activates the expression of multiple PRV genes and initiate the replication of PRV, which in turn travels across synapses to connected neurons. The PRV strain used in this study lacks the Us9 gene; therefore travels only in retrograde direction (Kratchmarov et al., 2013; Oyibo et al., 2014). The PRV^ΔIE^ carried EGFP, mCherry or Cre recombinase under the synapsin promoter.

We first tested the viral vectors with cultured 3T3 cells (a rodent cell line was chosen because PRV does not infect primate cells). Cells were infected with AAV-tTA together with AAV-IEo or AAV expressing wild type IE180 (AAV-IE180^WT^) (Fig.1B). 24 hrs later, PRV^ΔIE^-EGFP was added to the culture medium. PRV^ΔIE^-EGFP was titered down to a level where no EGFP-positive cells were observed if PRV^ΔIE^-EGFP was the only virus used to infect the cells. The cells were imaged 3 days later. Addition of AAV-IEo, but not AAV-IE180^WT^, led to the appearance of EGFP-positive cells, indicating that IEo efficiently activated EGFP expression. EGFP-positive cells concentrated in clusters, suggesting that PRV replicated and spread to neighboring cells.

The sensitivity of the AAV-IEo to Dox was tested in 3T3 cells as well (Fig. 1C). No EGFP-positive cells were observed when 1 μg/ml of Dox was added to the culture medium 1 hr before PRV^ΔIE^-EGFP infection, suggesting that Dox can completely inhibit PRV^ΔIE^-EGFP gene expression and replication. Similarly, AAV-rtTA and AAV-IEo combination led to the formation of EGFP-positive cell clusters only in the presence of Dox (Fig.1D).

This system was then tested *in vivo* with the dentate gyrus (DG)-CA3-septum circuit, where the granule cells in DG project to CA3 pyramidal cells, and the CA3 pyramidal cells further project to lateral septum. We first examined the transport of a replication-capable PRV expressing EGFP (PRV154) in this pathway (Fig. 1E). PRV-EGFP was injected to the septum unilaterally. The brains were fixed and sectioned 24, 36 or 55 hrs after the injection, respectively. At 24 hr, EGFP-positive cells were mainly confined to the injection site with sparse distribution at CA3. At 36 hrs and 55 hrs, EGFP-positive cells appeared successively at bilateral CA3 and DG, confirming the bi-synaptic pathway from DG to CA3 and further to septum. This demonstrates that PRV can be effectively retrogradely transported in this pathway. Next, we examined the transport of PRV^ΔIE^ in this circuit (Fig. 1F). We used PRV^ΔIE^-Cre in tdTomato reporter mice (Ai9 mice) to obtain greater sensitivity. PRV^ΔIE^-Cre was injected into septum unilaterally. Two months later tdTomato-positive neurons were observed at bilateral CA3 regions but not at DG, indicating that PRV^ΔIE^-Cre can be taken-up by axonal terminals at the septum and retrogradely transported to the soma at CA3; but without replication, no transsynaptic transport of PRV^ΔIE^ to DG occurred, consistent with previous observation (Oyibo et al., 2014).

We then tested if AAV vectors could restore PRV^ΔIE^ replication (Fig.1G). AAV-tTA and AAV-IEo were co-injected into CA3 unilaterally. 2 weeks later PRV^ΔIE^-Cre was injected to septum ipsilateral to CA3 injection. The brains were fixed and sectioned 3 weeks after PRV^ΔIE^ injection. Mice drank water containing Dox throughout the experiment starting from AAV injection till perfusion except during the 72 hours between 48 hours before and 24 hours after PRV^ΔIE^ injection. tdTomato-positive neurons were observed at DG ipsilateral to CA3 injection, but not at contralateral DG, indicating that the AAVs effectively activated the replication and retrograde transneuronal transport of PRV^ΔIE^-Cre from CA3 to the ipsilateral DG.

### 2) Temporally restricted replication minimized PRV neuronal toxicity

To determine if PRV^ΔIE^ produced neuronal toxicity *in vivo*, we compared the electrophysiological properties of infected neurons to that of non-infected control neurons (Fig. 2), We used AAV-rtTA to precisely time viral replication. AAV-rtTA and AAV-IEo were co-injected to CA3 of Ai9 mice. Two weeks later PRV^ΔIE^-Cre was injected to the contralateral CA1. Dox was provided 48 hours before to 24 hours after PRV injection. Three weeks later, acute brain slices containing DG were prepared and analyzed with whole-cell patch clamp recording. tdTomato-positive neurons (infected) and their neighboring non-fluorescent neurons (control) demonstrated similar resting membrane potentials, spontaneous synaptic currents, current injection-triggered action potentials and other electrophysiological properties (Fig.2 C–E). We also took confocal images of brain sections from mice receiving the same viral injection procedures. Consistent with physiology results, the morphology of the tdTomato-positive neurons appeared normal (Fig. 2B). Together, the results indicate that transient replication of PRV^ΔIE^ did not produce lasting cell toxicity.

### 3) Directed stepwise tracing of polysynaptic circuits by inducible reconstitution of PRV^ΔIE^

In the above tracing from septum to DG, PRV traveled along three brain regions but was only transported across one order of synapses. We went to test if IEo-mediated reconstitution of PRV^ΔIE^ could be applied multiple times along the circuit to track polysynaptic connectivity (Fig.3). We chose a circuit consisting of four brain regions: from DG to retrosplenial cortex (RSP) via CA3 and CA1. To examine the direction of synaptic projections along this circuit we made SynaptoTAG2 AAV, which was adapted from SynaptoTAG AAV (Xu and Sudhof, 2013). It contains EGFP fused to synaptobrevin 2, which labels synaptic terminals. P2A sequence replaces the internal ribosome entry site (IRES) in the original version to enhance the expression of Syb2-EGFP fusion protein; and tdTomato replaces mCherry to fill neuronal soma and axons as it is less prone to aggregation (Fig. 3A). As shown in Fig. 3B–D, SynaptoTAG2 AAV demonstrates the synaptic projections from DG to ipsilateral CA3, from CA3 to bilateral CA1, and from CA1 to ipsilateral RSP.

**Figure 3.**
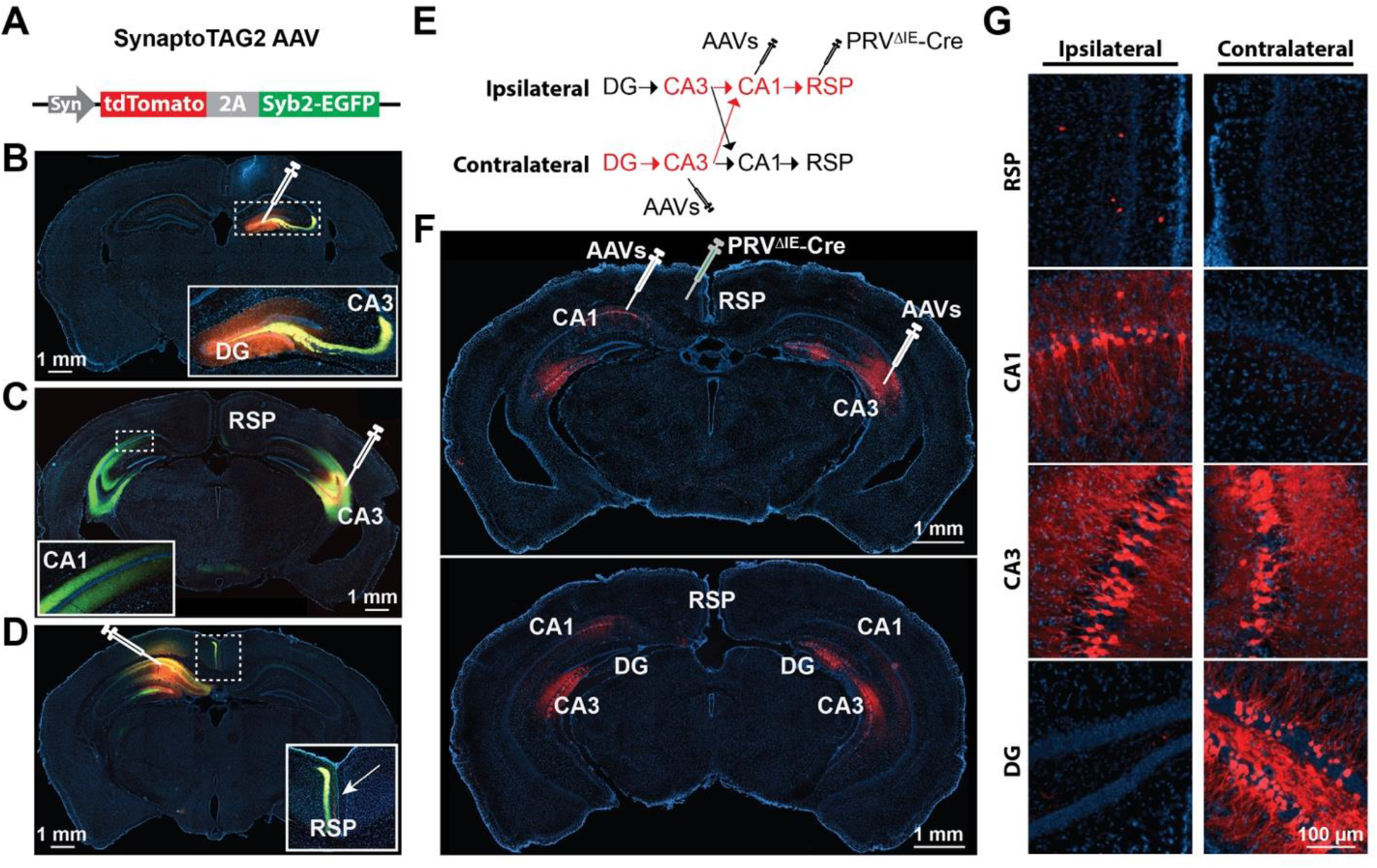
Controlled stepwise tracing of polysynaptic circuits by inducible reconstitution of PRV replication. **(A-D)** Tracing the synaptic pathways from DG to the retrosplenial area (RSP) with SynaptoTAG2 AAV. (**A**) SynaptoTAG2 AAV mediates bicistronic expression of tdTomato (labeling the soma and axons) and EGFP fused to synaptobrevin 2 (Syb2) (labeling synaptic terminals). (**B**) DG neurons project to ipsilateral CA3. The insert shows the enlargement of the area marked by dashed lines. (**C**) Neurons in CA3 project to bilateral CA1. (**D**) Neurons in CA1 project unilaterally to RSP. The arrow points to the midline of the brain. (**E**, **F**, **G**) Directed polysynaptic tracing from DG to RSP with PRV^ΔIE^-Cre. (**E**) Schematic showing the injection sites of the AAVs (AAV-tTA and AAV-IEo) and PRV^ΔIE^-Cre in Ai9 mice. Arrows indicate the direction of synaptic projections. (**F**) Two representative brain sections at different anterior-posterior positions showing the directed transsynaptic transport of PRV^ΔIE^-Cre. tdTomato-positive neurons were detected in the injection site (RSP), ipsilateral CA1, bilateral CA3 and contralateral DG. (**G**) Enlargement of the indicated brain regions in **F** (lower panel). See also Figure S2.

We then co-injected AAV-tTA and AAV-IEo into CA1 and contralateral CA3 of Ai9 mice, followed by injecting PRV^ΔIE^-Cre into ipsilateral RSP two weeks later (Fig. 3E). The mice drank Dox water except in the time window of 48 hours before to 24 hours after PRV injection. As shown in Fig. 3F–G, tdTomato-positive neurons were observed in: 1) RSP, the injection site; 2) ipsilateral CA1, reflecting axonal uptake and unilateral retrograde transport from RSP; 3) bilateral CA3, due to bilateral transsynaptic transport of PRV^ΔIE^-Cre from CA1 to CA3; and 4) contralateral DG, due to unilateral transsynaptic transport of PRV^ΔIE^-Cre from CA3 to DG. Therefore, PRV crossed two orders of synaptic connections to reach four brain regions. These results along with Fig. 1G indicate that AAV-mediated reconstitution of PRV replication can be used to trace polysynaptic pathways in pre-determined directions and to pass a pre-determined number of orders of synaptic connections.

### 4) Limited hippocampus-striatum direct projection

After establishing the viral systems for controlled transneuronal tracing, we applied them to delineate the wiring between the hippocampus and the striatum, two complex brain structures with close functional interactions. We first examined the direct synaptic projection from the hippocampus to the striatum with SynatoTAG2 AAV. When SynatoTAG2 AAV was injected into the dorsal CA1 (Fig. 4A) or CA3 (Fig. 4B), no EGFP-positive synapses were detected in either dorsal striatum or NAc although tdTomato-positive axons and EGFP-positive synaptic terminals from the hippocampus could be detected in the neighboring septum (Fig. 4A–B). When SynatoTAG2 AAV was injected into the ventral CA1 (some neurons in adjacent ventral CA3 were also infected), EGFP-positive synaptic terminals were observed in NAc with a concentration in NAc shell (Fig. 4C). The results show that the CA1 or CA3 regions in the dorsal hippocampus does not project directly to either dorsal striatum or NAc, while ventral hippocampus projects directly to NAc but not the dorsal striatum. Consistent with the anterograde tracing with SynatoTAG2 AAV, when PRV^ΔIE^-Cre alone was injected into the dorsal striatum of Ai9 mice, no tdTomato-positive neurons were found at either the dorsal or ventral hippocampus (Fig.4D); but when PRV^ΔIE^-Cre alone was injected into NAc, tdTomato-positive neurons were found in the ventral but not the dorsal hippocampus (Fig.4E). These data are consistent with early tracing studies showing that ventral hippocampus and the dorsal subiculum but not dorsal CA1 projected to NAc (Brog et al., 1993; Groenewegen et al., 1982; Groenewegen et al., 1987; Kelley and Domesick, 1982; McGeorge and Faull, 1989). In a recent study with AAV local infection, dorsal CA1 was found to project to NAc (Trouche et al., 2019). But we did not observe these projections with either anterograde or retrograde tracing.

**Figure 4.**
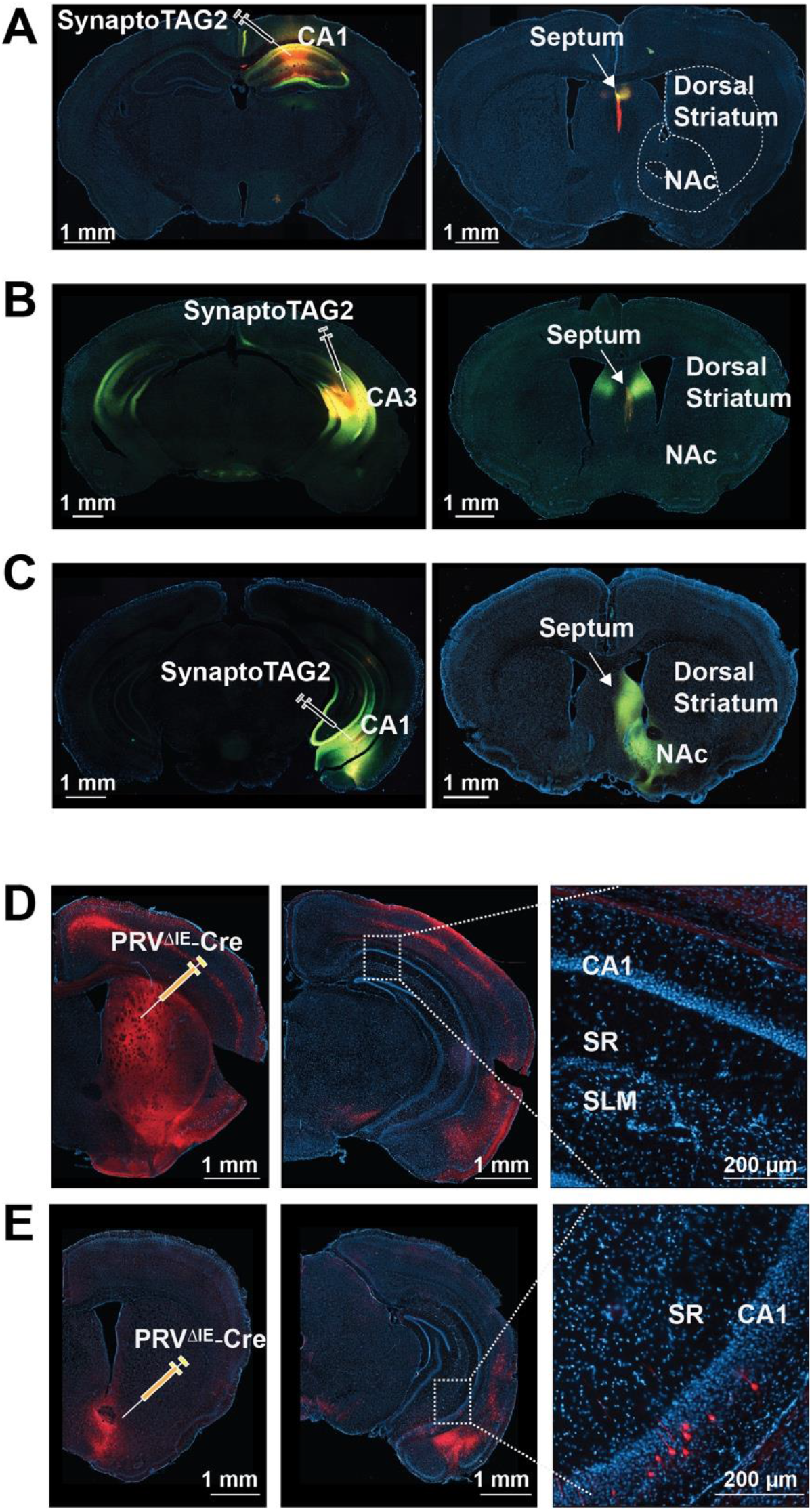
Limited hippocampus-striatum direct projection. (**A**) SynaptoTAG2 AAV was injected into the dorsal CA1. No EGFP-positive synaptic terminal were observed in the dorsal striatum or the nucleus accumbens (NAc). (**B**) SynaptoTAG2 AAV was injected into the ventral CA1 (part of the ventral CA3 was also infected). EGFP-positive terminals were distributed in NAc but not dorsal striatum. (**C**) SynaptoTAG2 AAV was injected into the CA3. No EGFP-positive synaptic terminals were observed in the dorsal striatum or the nucleus accumbens (NAc). (**D**) Ai9 mice received PRV^ΔIE^-Cre at the dorsal striatum. No tdTomato-positive neurons were observed in the dorsal or ventral hippocampus. (**E**) Ai9 mice received PRV^ΔIE^-Cre at NAc. tdTomato-positive neurons were observed in the ventral but not the dorsal hippocampus.

### 5) Directed tracing of disynaptic circuits between the CA1 and the dorsal striatum

To systematically trace the hippocampus-dorsal striatum pathways with the PRV^ΔIE^ system we picked 6 brain regions which receive direct hippocampal projections to determine if they wire the hippocampus to the striatum via polysynaptic connections (Fig. 5A). These candidate intermediate brain regions include the dSUB, PFC, lateral septum (LS), entorhinal cortex (EC), supramammillary nucleus (SUM) and RSP (Cenquizca and Swanson, 2007). When AAV-rtTA and AAV-IEo were co-injected into one of the candidate intermediate brain regions, followed by an injection of PRV^ΔIE^-Cre into the dorsal striatum two weeks later, distinct neuronal populations in the dorsal, intermediate, or ventral hippocampus expressed tdTomato (Fig. 5B). tdTomato was detected in the pyramidal cells and interneurons in the dorsal CA1 when PRV^ΔIE^-Cre passed dSUB. tdTomato-positive pyramidal cells were detected in the intermediate and ventral CA1 when AAVs were injected into the PFC, lateral septum and the EC. No tdTomato-positive pyramidal cells were detected in the hippocampus when AAVs were injected into the SUM or the rostral part of the RSP (rRSP) suggesting that they did not provide a bridge connecting the dorsal hippocampus to the dorsal striatum. When AAVs were injected into the dSUB, a significant number of tdTomato-positive pyramidal cells were detected at the CA3 and DG too, possibly because AAVs diffused to the CA1 and CA3 regions thus allowing PRV to cross synapses in the hippocampus.

**Figure 5.**
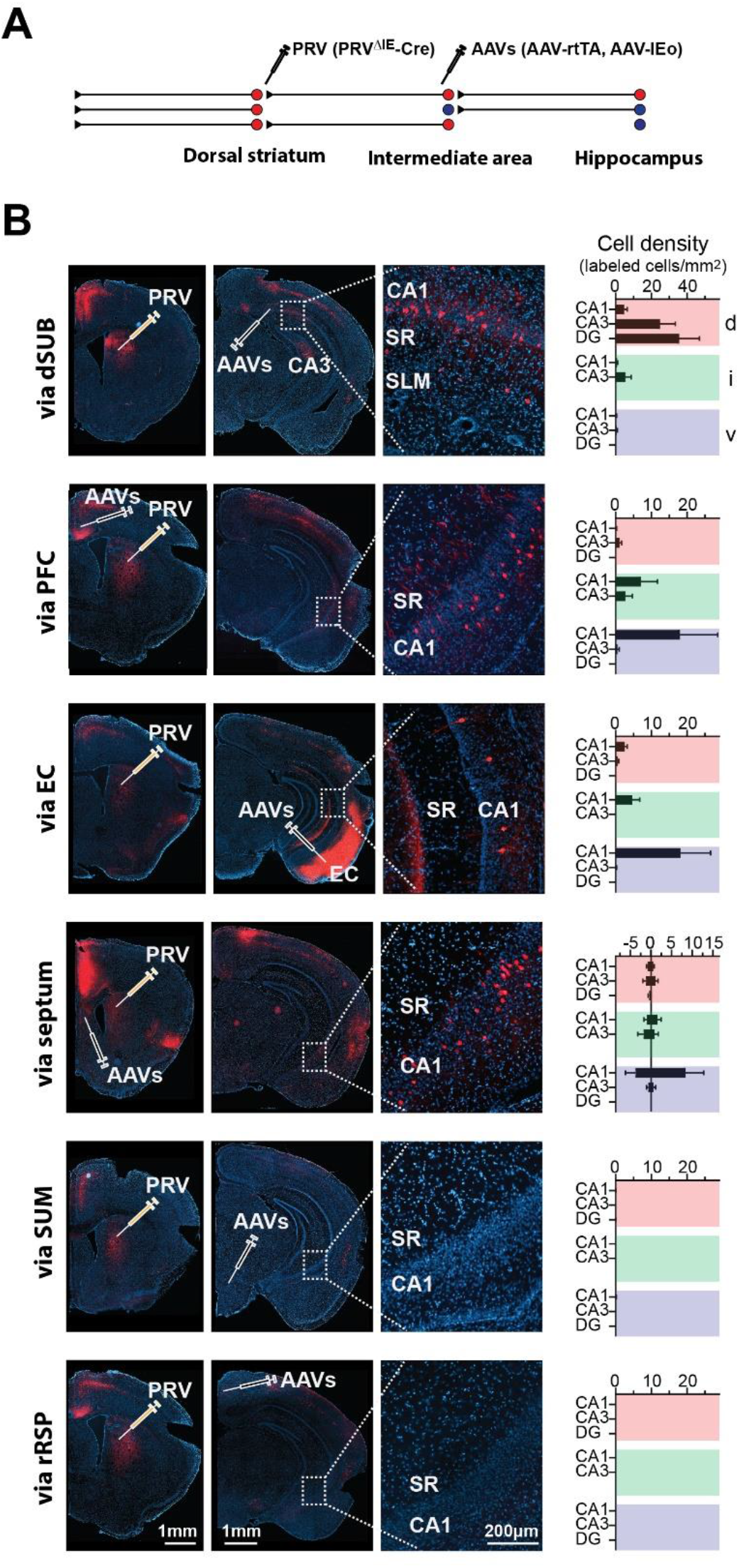
Tracing di-synaptic hippocampus-dorsal striatum circuits with PRV^ΔIE^. (**A**) Viral injection scheme for directed retrograde tracing of the hippocampus-dorsal striatum circuits. Ai9 mice received PRV^ΔIE^-Cre at the dorsal striatum and AAVs (AAV-rtTA and AAV-IEo) at a selected intermediate region. (**B**) Representative photos and quantification. tdTomato-positive neurons were detected in the dorsal CA1 when PRV^ΔIE^-Cre passed dSUB. PRV^ΔIE^-Cre also reached to the intermediate and ventral CA1 via the PFC, septum and the EC, respectively. Abbreviations: SR, stratum radiatum; SLM, stratum lacunosum-moleculare. Data are represented as mean ± SEM. n=17-33 brain sections from 3-4 mice for each intermediate region. In the bar graphs, positive values indicate that the cells were counted in the hemisphere ipsilateral to the viral injection sides; while the negative values mean that the cells were counted at the hemisphere contralateral to viral injection sides. tdTomato-positive cells were observed at the contralateral side only when AAVs were injected to the septum. The red, green and violet colors denote the dorsal (d), intermediate (i) and ventral hippocampus (v), respectively. Dorsal and intermediate hippocampus are demarcated at a horizontal line at the level of the ventral edge of the lateral blade of the dentate gyrus. The intermediate-ventral hippocampus border was a horizontal line at the level of the dorsal edge of the rhinal fissure.

### 6) Controlled polysynaptic tracing of the hippocampus-striatum-SNr circuits

We further extended the tracing of the hippocampus-striatum connectivity to SNr, a downstream target of the striatum. The dorsal striatum has two major populations of medium spiny neurons (MSNs)--those expressing dopamine D1 receptor, and those expressing dopamine D2 receptor. D1-positive MSNs primarily project to pars reticulata of SNr and compose the “direct pathway”, while D2-postive MSNs primarily project to the globus pallidus and form the “indirect pathway” (Calabresi et al., 2014). The direct and indirect pathways play opposing roles in the regulation of locomotion and reinforcement learning (Cui et al., 2013; Kravitz et al., 2012). Here we started the tracing at SNr by injecting PRV^ΔIE^-Cre into SNr. The AAVs for inducible IEo expression were injected into both dorsal striatum and the PFC (Fig. 6A). tdTomato-positive neurons were detected at SNr, the striatum, the PFC and ventral CA1 (Fig.6B). The results indicate that the hippocampus, via the PFC, can innervate the striatal direct pathway. It further demonstrates the effectiveness of this system in elucidating polysynaptic organization of neuronal circuits.

**Figure 6.**
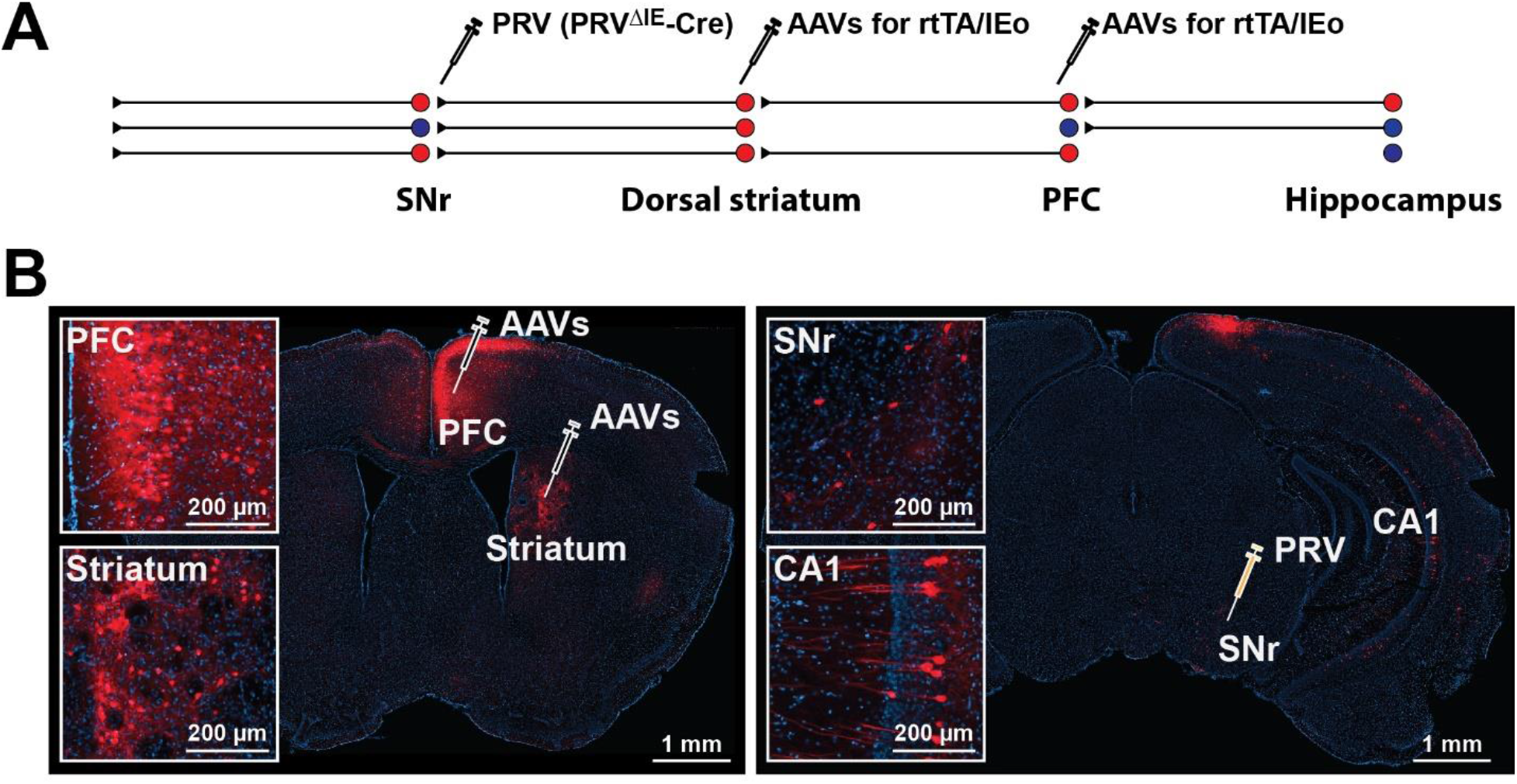
Tracing tri-synaptic hippocampus-striatum-SNr circuits with PRV^ΔIE^. (**A**) Viral injection scheme for retrograde tracing from the SNr to the hippocampus. Ai9 mice received PRV^ΔIE^-Cre at the SNr and the AAVs (AAV-rtTA and AAV-IEo) at the dorsal striatum and PFC. (**B**) Representative photos showing tdTomato-positive neurons in the SNr (the PRV^ΔIE^-Cre injection site), the dorsal striatum (direct presynaptic neuron to SNr neurons), the PFC (the intermediate region) and the CA1 of the hippocampus.

### 7) Directed tracing of disynaptic circuits between the CA1 and NAc

Similar to the experiments described above for dorsal striatum, we tested the 6 brain regions as candidate intermediate brain regions connecting the broad hippocampal regions to the NAc. AAV-rtTA and AAV-IEo were co-injected to one of the candidate intermediate brain regions of Ai9 mice, followed by an injection of PRV^ΔIE^-Cre in the NAc two weeks later (Fig. 7A). tdTomato was detected in the pyramidal cells and interneurons in the dorsal CA1 when AAVs were injected to dSUB. tdTomato-positive pyramidal neurons were detected at the dorsal CA1 and CA3 when AAVs were injected to lateral septum. PRV^ΔIE^-Cre traveled to the intermediate and ventral CA1 via the PFC and the EC. When the AAVs were injected into the SUM and rRSP, PRV^ΔIE^-Cre did not produce tdTomato-positive neurons in the hippocampal areas outside the ventral hippocampus directly projecting to NAc, suggesting that SUM and rRSP did not connect the hippocampus with NAc (Fig. 7B).

**Figure 7.**
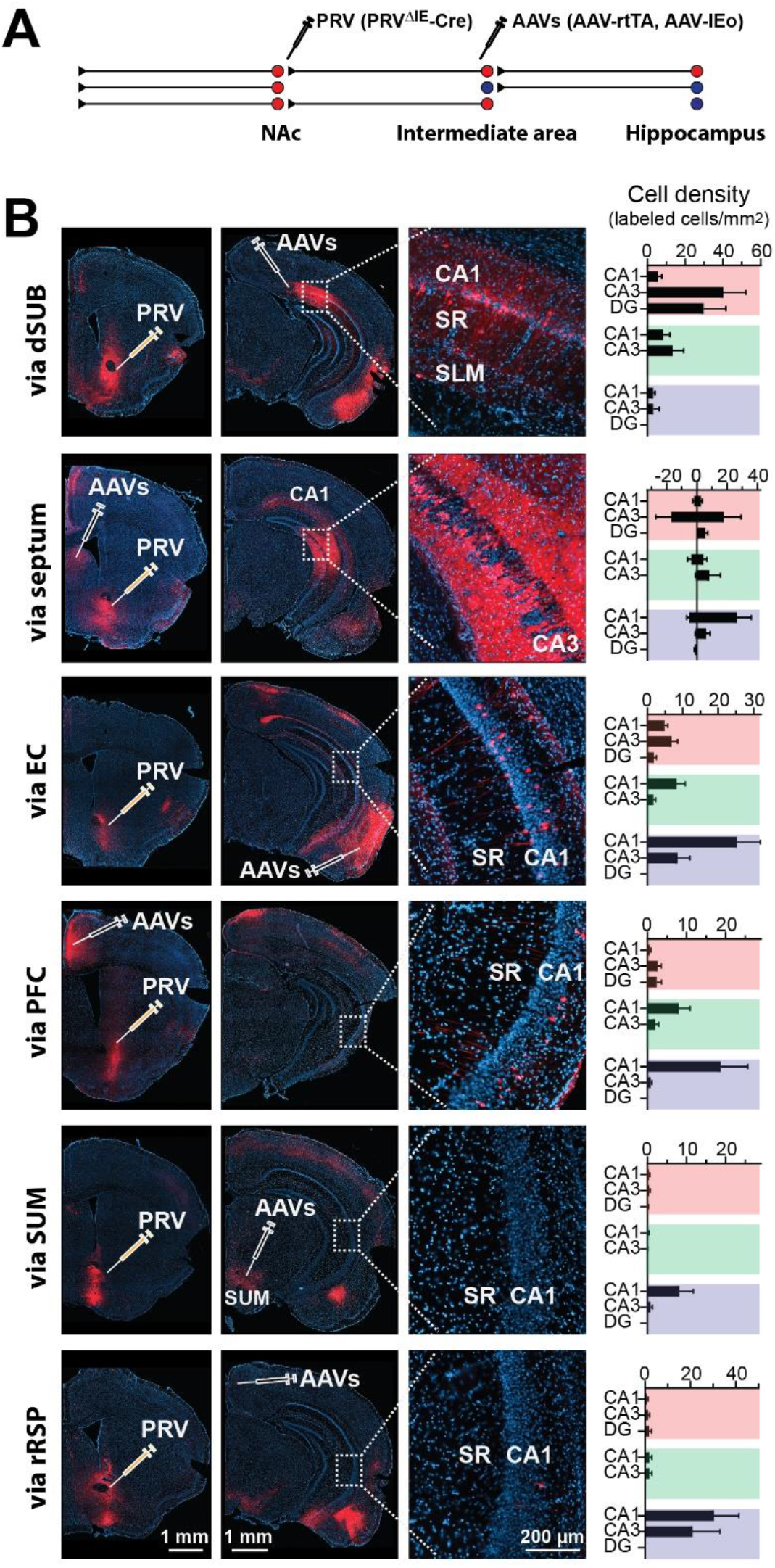
Tracing di-synaptic hippocampus-NAc circuits with PRV^ΔIE^. (**A**) Viral injection scheme showing the directed retrograde tracing of the hippocampus-NAc circuits with PRV^ΔIE^-Cre in Ai9 mice. The mice received PRV^ΔIE^-Cre at the NAc and AAVs (AAV-rtTA and AAV-IEo) in the indicated intermediate regions, respectively. (**B**) Representative photos and quantification showing that tdTomato-positive neurons were detected in different hippocampal regions when AAVs were injected into different intermediate regions. Data are represented as mean ± SEM. n=29-51 brain sections from 4-6 mice for each intermediate region. In the bar graphs, positive values indicate that the cells were counted in the hemisphere ipsilateral to the viral injection sides; while the negative values mean that the cells were counted at the hemisphere contralateral to viral injection sides.

Taken together, the hippocampal projections to the striatum (dorsal striatum and NAc) demonstrate the following connectivity patterns (summarized in Fig.8). (1) The dSUB is the major passageway for the dorsal CA1 to connect to the dorsal striatum and the NAc. (2) The EC and PFC are the major transfer stations connecting ventral CA1 to dorsal striatum and NAc. (3) The rostral part of RSP does not connect the hippocampus to the striatum although it receives dense inputs from CA1 (Fig. 3D). (4) Dorsal CA1 send dense inputs to the SUM (Cenquizca and Swanson, 2007) and the SUM neurons project to the NAc (Fig. 7B, SUM panel) (Vertes, 1992), but SUM does not connect the dorsal CA1 to the NAc. (5) Lateral septum (LS) primarily connects the dorsal CA1 and CA3 to the NAc, and connects the ventral CA1 to the dorsal striatum. (6) Most connectivity is unilateral except that LS connects the striatum to the hippocampus in both hemispheres. (7) Tracing of the hippocampus-SNr polysynaptic pathway indicates that the dorsal striatum direct pathway can be innervated by the hippocampus through CA1-PFC projections.

**Figure 8.**
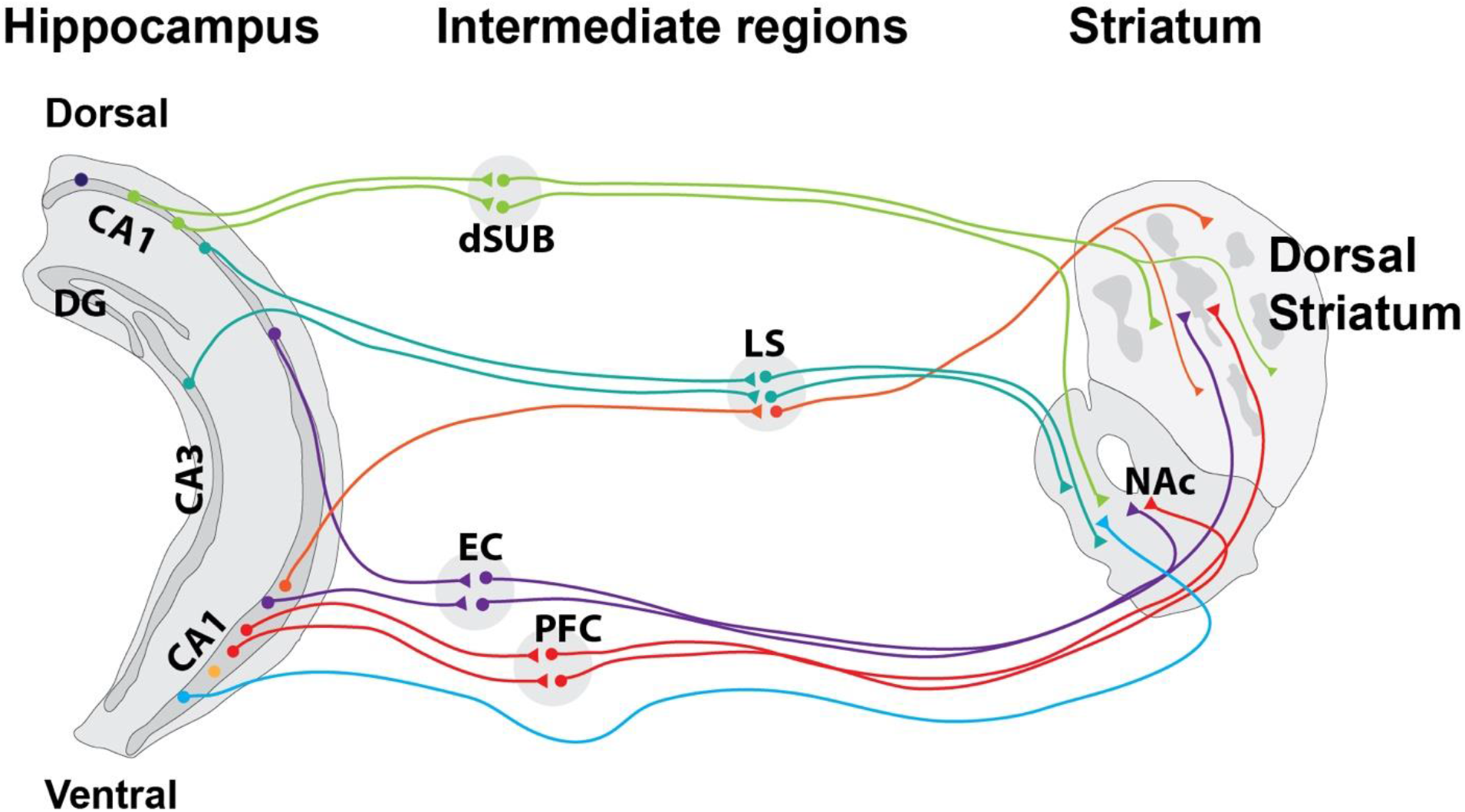
Hippocampus-striatum wiring diagram. Schematic summarizing the connectivity revealed in Fig. 4–7.

## Discussion

### Controlled polysynaptic tracing of neuronal circuits

Modern neuroscience started over a century ago when Santiago Ramon Cajal’s study of brain histology led to a connectionist view of the neuronal systems: the brain was a network of diverse neuronal cells (Swanson and Lichtman, 2016). Since then extensive research in neuronal wiring has produced important insights into brain structural organization. A large collection of neuronal tracers has been developed. For example, exogenous fluorescent proteins or other markers have recently been widely used to trace the projections of specific types of neurons (Bienkowski et al., 2018; Livet et al., 2007; Viswanathan et al., 2015). Transneuronal viruses, originally used as self-amplifying tracers, have been modified for monosynaptic tracing. In a pioneering work, glycoprotein was removed from recombinant RV to prevent RV from spreading across synapses (Wickersham et al., 2007a). Transneuronal spreading of RV can then be reconstituted by expressing the glycoprotein in-trans. This allows monosynaptic inputs to be traced from “starter” neurons that express the glycoprotein (Wickersham et al., 2007b). Similar strategies were used to generate recombinant PRV (Kondoh et al., 2016; Oyibo et al., 2014; Wu et al., 2014), VSV (Beier et al., 2013), and HSV (Lo and Anderson, 2011; Zeng et al., 2017). This strategy can be combined with other tools to trace circuits spanning three brain regions albeit the viruses pass only one order of synaptic connection (Schwarz et al., 2015; Xu et al., 2016).

However, we still know very little about the long-range polysynaptic circuits and need technology to probe the polysynaptic neuronal organization. In the current study, we demonstrate that inducible, directed, stepwise tracing of the polysynaptic circuits can be realized with transcomplementation of PRV. This work is built on a long history of engineering PRV vectors. PRV is one of the first viruses used for tracing (Card et al., 1991; Ekstrand et al., 2008) and has been modified to trace from specific neuronal types for a variety of applications (DeFalco et al., 2001; Pomeranz et al., 2017). The recent finding that PRV lacking IE180 showed no cytotoxicity opened up many opportunities (Oyibo et al., 2014; Wu et al., 2014). Here we found that codon-optimized IE180--IEo--significantly improved the efficiency of transcomplementation *in vitro* and *in vivo*, and that inducible expression of IEo minimized neuronal toxicity. With this system, viral spreading can be directed to a specific pathway without affecting unintended pathways. The number of orders of synaptic connections to be traced can be pre-determined by controlling where exactly IEo is expressed. Furthermore, this system can carry recombinase or other effectors to control or analyze the functions of a specific pathway. Importantly, although the current study focuses on the hippocampus-striatum wiring, the same strategy can be extended to unravelling other brain circuits.

While being a powerful tool, this PRV system can be improved further in the future so that it can be used more conveniently. For example, each brain region is made up of heterogeneous neurons (Zeng and Sanes, 2017). The IEo expression can also be made conditional, so that we can examine the roles of specific neuronal types in the circuits. With the current PRV system, we cannot exclude the possibility that PRV replicates and spreads via local interneurons in each of the intermediate regions, but if the replication is restricted to a specific type of neurons, this possibility can be minimized. In addition, IE180 gene needs to be turned on before or at the time of PRV injection to enable transneuronal spreading, possibly because PRV becomes dormant in the absence of IE180. Therefore, the AAVs encoding IE180 need to be injected to all the brain regions of interest before PRV injection. Based on the findings of the molecular mechanism for PRV latency, we can develop a way to reactivate PRV replication and spreading (for instance, applying deacetylase inhibitors), which may provide a more flexible and convenient system.

### Hippocampus-striatum polysynaptic wiring diagram

The positioning and functions of the hippocampus and the striatum in the brain systems suggest that there must be an extensive interaction between the two in multiple brain functions. The hippocampus is located at the top of the hierarchy of cortical information processing (Squire et al., 2004). It is particularly important in integrating multimodal sensory information into representations of the context (Maren et al., 2013); and in encoding and recording the temporal or spatial relationship among events or objects (Eichenbaum, 2017), which may also include sequence information (Davachi and DuBrow, 2015). The striatum is an interface between the limbic system and motor systems (Pennartz et al., 2011). It plays a particular important role in mediating memory-guided behaviors and motor sequence learning. The positioning of the hippocampus and the striatum in the brain systems suggests that there must be an extensive interaction between the two in multiple brain functions.

The hippocampus has a prominent dorsal-ventral functional division. The hippocampus is an elongated and laminated structure. Along its long (dorsal-ventral) axis, each layer contains DG, CA3, CA2 and CA1 subregions (van Strien et al., 2009) (Bienkowski et al., 2018). These subregions compute information differentially (Tonegawa and McHugh, 2008). A functional division has also been noticed long ago between the dorsal and ventral hippocampus (corresponding to the posterior and anterior hippocampus in human) (Fanselow and Dong, 2010; Moser et al., 1993). The dorsal hippocampus is preferentially engaged in cognitive processing of memory-related neuronal information, while the ventral hippocampus is more involved in emotional information processing (Fanselow and Dong, 2010). The functional division of the dorsal and ventral hippocampus is well underpinned by the distinct neuronal gene expression profiles and neuronal connectivity of these regions. For example, based on the expression of a large collection of genes, the hippocampus can be divided into dorsal, intermediate and ventral hippocampus (Dong et al., 2009). The different gene expression levels contribute to the functional heterogeneity. For example, the gradient of HCN1 expression along the dorsal-ventral axis may directly determine the size of place field of place cells (Hussaini et al., 2011). More importantly, the dorsal and ventral hippocampus differ in synaptic connectivity. In terms of inputs, the entorhinal inputs to the hippocampus are topographically arranged along hippocampal long axis; the amygdala, the thalamus (especially nucleus reuniens) and the hypothalamus preferentially project to the ventral hippocampus. In terms of the synaptic outputs, the dorsal vs. ventral hippocampal projections to the lateral septum, amygdala and the NAc are all topographically arranged (Strange et al., 2014).

Similar to the hippocampus, the striatum has a prominent dorsal-ventral functional division. The striatum is traditionally divided into the dorsal striatum and NAc. The dorsal striatum is further divided into dorsolateral and dorsomedial striatum. The three compartments--dorsolateral, dorsomedial striatum and NAc--are arranged largely from dorsal to ventral direction along an axis around 45 degrees to the brain midline (Voorn et al., 2004). The three compartments show apparent functional division. The dorsolateral striatum is preferentially engaged in procedure learning and stimulus-response learning; the dorsomedial striatum is involved in spatial learning; and NAc is particularly involved in reinforcement learning. Learning and habit formation requires a dynamic interaction among the ventral, dorsomedial and dorsolateral striatum (Thorn et al., 2010) (Belin and Everitt, 2008). Similar to the hippocampus, the dorsal-ventral gradient of the striatum is manifested by differential gene expression patterns, but more prominently by the distinct synaptic connectivity (Voorn et al., 2004). The dorsolateral striatum is innervated by excitatory inputs from motor and somatosensory cortices; the dorsomedial striatum receives strong inputs from the medial prefrontal cortex and the thalamus; and NAc receive strong direct inputs from the prefrontal cortex, amygdala, and the ventral hippocampus.

Although the hippocampus and the striatum have significant functional interactions as mentioned above, earlier studies have focused on the collaboration between the ventral hippocampus and NAc, likely due to direct connections between them (Pennartz et al., 2011). But the functional interaction can be well mediated through polysynaptic pathways. Here with the new PRV^ΔIE^ system, we delineated the polysynaptic wiring diagram from the hippocampus to striatum, which opens the door to more extensive study of the hippocampus-striatum interactions. The results demonstrate that specific anatomical domains in the hippocampus are connected to specific anatomical domains in the striatum via distinct intermediate brain regions. These separate tracts may organize neurons in multiple regions into distinct functional channels, and each channel may selectively process one type of information. Particularly, our tracing reveals a few pathways linking the dorsal hippocampus to multiple regions of the striatum. Similarly, the tracing also indicates that the lateral septum links the ventral hippocampus to the dorsal striatum. Considering the roles of the dorsal hippocampus in spatial representation and declarative memory, it is not surprising that these pathways will be critically involved in these brain functions. Since PRV^ΔIE^-Cre system can turn on or off the expression of genes, we can use it to express in the selected pathways specific tools for functional manipulations, such as optogenetics or chemogenetic tools, and so on. These manipulations will allow us to determine the functions of the specific functional channel between the hippocampus and the striatum.

## Acknowledgments

We thank Dr. Ege Kavalali for critical comments and suggestions on the manuscript, the laboratories of Dr. Lynn W. Enquist (Princeton University) and Dr. Anthony Zador (Cold Spring Harbor Laboratory) for the replication-deficient PRVs expressing Cre, mCherry or EGFP, the CNNV core for PRV154. This study was supported by Klingenstein-Simons Fellowship Awards in Neuroscience (to WX) and grants from NIH/NINDS (NS104828). The UT BRAIN seed grants (365231) and the Texas Institute for Brain Injury and Repair (TIBIR) pilot grant provided funds to initiate this study. We also thank Dr. Denise Ramirez and Dr. Julian Meeks (the Whole Brain Microscopy Facility at UT Southwestern) for help with scanning the whole-mount brain sections and Dr. Shin Yamazaki (UT Southwestern) for help with confocal imaging.

